# Anti-Microbial Resistance in Agents Causing Urinary Tract Infections

**DOI:** 10.1101/316869

**Authors:** Zuhair Ali Rizvi, Ali Murad Jamal, Ali Hassan Malik, Naimat Ullah, Daneyal Arshad

## Abstract

**BACKGROUND:** Urinary Tract Infections are usually treated with empirical therapy by physicians based on previous knowledge of predictability of causative agents and their antimicrobial susceptibilities.

**OBJECTIVE:** The objective of this study was to determine the frequency of various pathogens causing urinary tract infections and their antimicrobial susceptibility in patients presenting in out-patient department of a tertiary care hospital.

**MATERIALS AND METHODS:** This descriptive cross sectional study was conducted in Out-Patient department of Urology of Benazir Bhutto Hospital during a period of 6 months from January 2017 to June 2017 after ethical approval from institutional research forum of Rawalpindi Medical University. 1000 patients (12 years old or above) that were clinically suspected for urinary tract infections were included in this study. Patients with co-morbidities like diabetes mellitus, renal pathologies, Immunodeficiency disorders, malignancies and congenital urogenital disorders were also excluded. Recipients of corticosteroid therapy or with a history of intake of broad spectrum antibiotics in previous 15 days were also excluded. Modified Kirby-Bauer disc diffusion method was used for determining the antimicrobial resistance against various antimicrobials.

**RESULTS:** A total of 530 (53%) isolates were found to be culture positive for E.coli(77.4%),Klebsiella (6.4%), Enterobacter (6.0%), Pseudomonas (3.8%), Staphylococcus saprophyticus (3.4%), Citrobacter (1.1%) and Morganella (0.4%).

Antimicrobial resistance against commonly used antimicrobials was alarmingly high

**CONCLUSION:** Surveillance of trends of antimicrobial susceptibility pattern is highly important.

## INTRODUCTION

Urinary Tract Infections (UTI) are one of the most commonly diagnosed diseases in Out-Patient Department. (1) The selection of antibiotic therapy by a physician is based on knowledge of prevalent microorganisms in the setting and recent updates about the antimicrobial susceptibility patterns and the clinical status of the patient. (2)

Although E. coli is the most commonly isolated microorganisms from patients with UTI, however a significant difference in the prevalence has been stated. The percentage isolation of E. coli varied from 26.1% in a Turkish study to 55.1% in a developing country like India. (3, 4)

Patients with diabetes are at four times more risk of developing Urinary Tract as compared to non-diabetics (5)

High resistance of agents causing UTI against commonly used antimicrobials has been reported. In a study conducted in Ethiopia, it has been stated that 93.3%, of the isolates were sensitive to gentamicin. Similarly, 60%, 60%, 56.6%, 46.6%, 40%, 33.3%, of the isolates were sensitive to chloramphenicol, nitrofurantoin, ciprofloxacin, Trimethprime-Sulfamethoxazole (TMP-SMX), ceftriaxone and nalidixic acid, respectively. (6)

Therefore, there was a need to conduct a study to determine most common agents associated with Urinary Tract Infections and their current antimicrobial resistance patterns so as to formulate a better antimicrobial therapy.

The objective of this study was to determine frequency of Urinary Tract Infections, agents causing UTI and their current antimicrobial resistance profiles.

## MATERIALS & METHODS

This descriptive cross sectional study was conducted among patients presenting in Out-Patient Department of Urology of Benazir Bhutto Hospital, Rawalpindi, after ethical approval from Institutional Research Forum of Rawalpindi Medical University.

Patients above the age of 12, presenting in above stated out-patient departments and clinically suspected for Urinary Tract Infections (UTI) were included in this study.

Patients who had no symptoms related to UTI were excluded. Patients with co-morbidities like diabetes mellitus, renal pathologies, Immunodeficiency disorders, malignancies and congenital urogenital disorders were also excluded on the basis of history. Recipients of corticosteroid therapy or with a history of intake of broad spectrum antibiotics in previous 15 days were also excluded.

Non cooperative female patients who didn’t give consent or refused to provide necessary information were also excluded.

The subjects included in the study were provided with a wide-mouthed standard sized sterile container. They were advised to clean the area around urethra with water before collecting the sample, let the area dry and collect the sample by catching the mid-stream with container being held at 2-3 inches away.

50 μl of uncentrifuged urine was taken on a clean slide and a cover slip was placed on it. The slide was then viewed under microscope. Presence of blood cells, epithelial cells, pus cells or cast bodies was duly noted. The presence of ten or more pus cells per high power field was considered as significant pyuria.

Gram staining technique was employed. Detection of at least one or more morphologically similar bacteria per oil immersion field was called significant.

10μL of specimen was transferred to MacConkey’s agar plate using a calibrated loop method. The agar plates were incubated at 35-37°C for 24 hours for identification of lactose fermenting and non-lactose fermenting bacteria. A specimen was considered positive for UTI if a single organism was cultured at a concentration of 10^5^ Colony Forming Units/ml.

CLED (Cystein Lactose Electrolyte Deficient) medium was further used for identification and isolation of urinary pathogens.

Modified Kirby-Bauer disc diffusion method was used for determining the antimicrobial susceptibility. The colonies were placed on afar plates using sterile inoculating wire loop. Antibiotic disks were placed using sterile forcep. The plates were left for 1 hour at room temperature to allow diffusion of antibiotics from the disks. The agar plates were again incubated for 24 hours at 37°C. CLSI guidelines were used to determine antimicrobial susceptibility.

Antimicrobial susceptibility and resistance was tested against Ampicillin, Amoxicillin + Clavulanic Acid (AMC), Gentamycin, Amikacin, Cefoperazone, Ceftazidime, Cefixime, Ceftriaxone, Cefepime, Piperacillin+Tazobactam (TZP), Cefoparazone+Sulbactam (CFP+SUL), Carbepenam, Fosfomycin, Trimethoprim/sulfamethoxazole (SXT), Ciprofloxacin, Ofloxacin, Levofloxacin, Norfloxacin and Nitrofurantoin. Data was analyzed using SPSS v22.0 and descriptive statistics were applied.

Data was entered and analyzed using SPSS v22.0. Following Operational Definitions were used:

- Microscopy findings of more than 10 WBC per high power field were called significant pyuria.
- Significant bacteriuria was defined as culture of a single bacterial species from the urine sample at a concentration of more than 100,000 cfu/ml.

## RESULTS

Out of 1000 patients, 303 (30.3%) males and 697 (69.7%) females, who were clinically suspected for Urinary Tract Infection, 530 (53%) patients, 160 (52.8%) males and 370 (53.0%) females were found to be culture positive while 430 (43.0%) patients, 143 (47.2%) males and 327 (47%) females, were culture negative.

Of 530 culture positive specimens, E.coli was most frequently isolated followed by *Klebsiella spp., Enterobacter spp., Pseudomonas spp., Staphylococcus spp., Proteus spp., Citrobacter spp.*, and *Morganella spp.*The difference in gender wise distribution of various pathogens was statistically significant p<0.05. (Table 1)

**Table 1:** Frequency of isolation of various microorganisms from urinary samples.

| ORGANISM | NUMBER | % AGE | MALE |  | FEMALE |  |
| --- | --- | --- | --- | --- | --- | --- |
|  |  |  | N | % | N | % |
| *E. coli* | 410 | 77.4 | 116 | 72.5 | 294 | 79.5 |
| *Klebsiella spp.* | 34 | 6.4 | 10 | 6.2 | 24 | 6.5 |
| *Enterobacter spp.* | 32 | 6 | 4 | 2.5 | 28 | 7.6 |
| *Pseudomonas spp.* | 20 | 3.8 | 14 | 8.8 | 6 | 1.6 |
| *Staphylococcus spp.* | 18 | 3.4 | 10 | 6.2 | 8 | 2.2 |
| *Proteus spp.* | 8 | 1.5 | 4 | 2.5 | 4 | 1.1 |
| *Citrobacter spp.* | 6 | 1.1 | 2 | 1.2 | 4 | 1.1 |
| *Morganella spp.* | 2 | 0.4 | 0 | 0.0 | 2 | 0.5 |

To test the null hypothesis that the “Distribution of Age is same across all categories of Organism”, Independent Samples Kruskal-Wallis test was applied. P<0.05 was considered significant. The null hypothesis was rejected (p=0.047). The distribution of age among various organisms was not same.

To compare mean age of isolation of various pathogens in males and females, independent sample “t” test was applied. P<0.05 was considered significant. The difference in means of isolation of various pathogens as per gender was significant for E. coli, *Enterobacter spp., Staphylococcus spp.* and *Citrobacter spp*. (Table 2)

**Table 2:** Mean age of distribution of various microorganisms in context with gender.

| ORGANISM | MEAN AGE | MALE | FEMALE | Independent sample “t” test |
| --- | --- | --- | --- | --- |
|  |  | N | N | P value |
| *E. coli* | 47.34 | 53.82 | 44.66 | $P=<0.0001$ |
| *Klebsiella spp.* | 42.65 | 52.00 | 38.75 | 0.0611 |
| *Enterobacter spp.* | 46.50 | 73.50 | 42.64 | 0.0056 |
| *Pseudomonas spp.* | 62.80 | 59.43 | 70.67 | 0.0913 |
| *Staphylococcus spp.* | 47.22 | 56.80 | 35.25 | 0.0040 |
| *Proteus spp.* | 40.75 | 31.50 | 50.00 | 0.2324 |
| *Citrobacter spp.* | 54.67 | 39.00 | 62.50 | 0.0004 |
| *Morganella spp.* | 43.00 | 43.00 | - | - |
| Independent Samples<br>Kruskal-Wallis test | $P = 0.047$ | | | |

Antimicrobial resistance of various pathogens against Ampicillin, Amoxicillin + Clavulanic Acid (AMC), Gentamycin, Amikacin, Cefoperazone, Ceftazidime, Cefixime, Ceftriaxone, Cefepime, Piperacillin+Tazobactam (TZP), Cefoparazone+Sulbactam (CFP+SUL), Carbepenam, Fosfomycin, Trimethoprim/sulfamethoxazole (SXT), Ciprofloxacin, Ofloxacin, Levofloxacin, Norfloxacin and Nitrofurantoin was determined. (Table 3)

**Table 3:** Anti-microbial resistance in various microorganisms against commonly used drugs.

| Antimicrobial Agent | *E. coli* | *Klebsiella spp.* | *Enterobacter spp.* | *Pseudomonas spp.* | *Staphylococcus spp.* | *Proteus spp.* | *Citrobacter spp.* | *Morganella spp.* |
| --- | --- | --- | --- | --- | --- | --- | --- | --- |
| Ampicillin | 87.3 | 100.0 | 50.0 | 0.0 | 100.0 | 100.0 | 100.0 | 100.0 |
| AMC | 62.7 | 100.0 | 50.0 | . | 0.0 | 2.0 | 100.0 | 100.0 |
| Gentamycin | 31.0 | 43.8 | 86.7 | 40.0 | 12.5 | 33.3 | 66.7 | 0.0 |
| Amikacin | 1.0 | 25.0 | 0.0 | 20.0 | 0.0 | 33.3 | 0.0 | 0.0 |
| Cefoparazone | 100.0 | 100.0 | 100.0 | 20.0 | . | 100.0 | 0.0 | 0.0 |
| Ceftazidime | . | . | . | 22.2 | . | 50.0 | 0.0 | . |
| Cefixime | 66.0 | 50.0 | 66.7 | 91.7 | . | 25.0 | 0.0 | 0.0 |
| Ceftriaxone | 63.7 | 47.1 | 66.7 | 100.0 | . | 25.0 | 0.0 | 0.0 |
| Cefepime | 63.2 | 43.8 | 66.7 | 22.2 | . | 0.0 | 0.0 | 0.0 |
| TZP | 11.3 | 17.6 | 33.3 | 22.2 | . | 0.0 | 66.7 | 0.0 |
| CFP+SUL | 10.3 | 29.4 | 33.3 | 22.2 | . | 0.0 | 66.7 | 0.0 |
| Carbepenam | 2.5 | 5.9 | 16.7 | 22.2 | . | 33.3 | 0.0 | 0.0 |
| Fosfomycin | 2.0 | 0.0 | 20.0 | . | 33.3 | 100.0 | 33.3 | 100.0 |
| SXT | 71.9 | 52.9 | 66.7 | . | 33.3 | 25.0 | 66.7 | 0.0 |
| Ciporfloxacin | 75.4 | 47.1 | 87.5 | 60.0 | . | . | 33.3 | 0.0 |
| Ofloxacin | 100.0 | . | . | 70.0 | . | . | 0.0 | . |
| Levofloxacin | 93.3 | 88.9 | 100.0 | 77.8 | 75.0 | 50.0 | . | . |
| Norfloxacin | 75.4 | 40.0 | 0.0 | 70.0 | 42.9 | 33.3 | 66.7 | 0.0 |
| Nitrofurantoin | 10.7 | 70.6 | 0.0 | 66.7 | 0.0 | 100.0 | 33.3 | 100.0 |

## DISCUSSION

In a study conducted in Iran, It has been stated that of all the urinary samples included in the study, 70.3% belonged to female patients while 29.7% belonged to male patients which is in accordance with our study. We observed that 69.7% belonged to female while 30.3% samples belonged to male patients. (1) It has also been stated by the same study that the difference between gender-wise distribution of various pathogens is statistically significant (p=<0.05) which is also in accordance with our study where p<0.05. (7)

Urinary Tract Infections are becoming difficult to treat owing to increasing resistance (8)

Emergence of resistance against broad spectrum antibiotic agents such as extended spectrum Beta Lactams, Fluoroquinolones and Carbepenams is causing problems worldwide. (9)

E.coli is the most commonly isolated pathogen from urinary samples. In a study conducted in United States of America, it was observed that 53.6% samples were culture positive for E. coli, 14.6% for Proteus, 13.9% for Klebsiella, 4.5% for Enterococcus and 4.1% for Staphylococcus. (10)

According to the same study, 60 % strains of E. coli were resistant to Fluoroquinlones, 7% to Nitrofurantoin, and 27% to SXT. (4) while according to our study, 75.4% to 100% strains of E. coli were resistant to Fluoroquinolones, 10.7% to Nitrofurantoin and 71.9% to SXT (10)

In another study, it was observed that 60% of isolates from urinary samples were of E.coli, 12% of Klebsiella and 8% for Enterococcus. (11)

In a study conducted in India, it was observed that frequency rate for E.coli was 69.8% followed by Klebsiella to be 7.9%, 4.8% for Staphylococcus, 4.8% for Pseudomonas, 4.8% for Enterococcus and 1.6% for MRSA. (12) These results were comparable to our observations

It was further stated that 81.8% strains of E.coli were resistant to Ampicillin, 72.7% to AMC, 54.5% to Cefepime, 21.5% to (CFP+SUL), 2.3% to 6.8% to Carbepenams, 36.4% to Gentamicin, 9.1% to Amikacin, 65.9% to Ciprofloxacin, 61.4% to Levofloxacin, 47.7% to SXT and 15.9% to Nitrofurantoin. These resistance rates were also comparable to our findings. (12)

According to a study, only 57% individuals out of those with provisional diagnosis of Urinary Tract Infections were culture positive for Uropathogens which is comparable to 53% in our study. (13)

According to a study conducted in Indonesia, it was observed that resistance of E. coli against Fluoroquinolones ranged from 41.4% for Levofloxacin to 71.6% for Ciprofloxacin while 78.4% isolates of Klebsiella were resistant to Levofloxacin. Less than 5% strains of E.coli and Less than 20% strains of Klebsiella were resistant to Fosfomycin.

It was observed that 85.5% strains of E.coli were found to be resistant to Ampicillin, 50.5% to AMC, 32.0% to Gentamicin, 58.1% to Ceftriaxone, 2.5% for Carbepenams, 0.9% to Fosfomycin and 13.1% to Nitrofurantoin while according to our study values were 87.3%, 62.7%, 31.0%, 63.7%, 2.5%, 2.0% and 10.7 % respectively.

It was also stated that 40.5% strains of Klebsiella were resistant to Gentamicin, 5.4% to Carbepenams, 78.4% to Levofloxacin, 83.8% to Nitrofurantoin while according to our observations, resistance rates were 43.8%, 5.9%, 88.9% and 70.6% respectively. (14)

Ilić et al has stated that frequency of E.coli was isolated from 67.7% samples and 69.5% of strains of E.coli were found to be resistant to Ampicillin which is in accordance with our study. (15)

An American study has stated that resistance among isolates of E.coli from urinary samples was lowest for Nitrofurantoin (<1%). But according to our observations, resistance rates of E.coli against Nitrofurantoin were found to be 10.7%. (16)

Our study is also in agreement with previously done studies which state that 61% strains of Klebsiella were resistant to Extended Spectrum Beta Lactams. (17)

In a few other studies, the resistance rates of E. coli against Nitrofurantoin have been stated to be less than 15% which is in accordance with our study. (18,19)

